# Distinct effects of different matrix proteoglycans on collagen fibrillogenesis and cell-mediated collagen reorganization

**DOI:** 10.1101/2020.08.30.274001

**Authors:** Dongning Chen, Lucas R. Smith, Gauri Khandekar, Pavan Patel, Christopher K. Yu, Kehan Zhang, Christopher S. Chen, Lin Han, Rebecca G. Wells

## Abstract

The extracellular matrix (ECM) is a complex mixture composed of fibrillar collagens as well as additional protein and carbohydrate components. Proteoglycans (PGs) contribute to the heterogeneity of the ECM and play an important role in its structure and function. While the small leucine rich proteoglycans (SLRPs), including decorin and lumican, have been studied extensively as mediators of collagen fibrillogenesis and organization, the function of large matrix PGs in collagen matrices is less well known. In this study, we showed that different matrix PGs have distinct roles in regulating collagen behaviors. We found that versican, a large chondroitin sulfate PG, promotes collagen fibrillogenesis in a turbidity assay and upregulates cell-mediated collagen compaction and reorganization, whereas aggrecan, a structurally-similar large PG, has different and often opposing effects on collagen. Compared to versican, decorin and lumican also have distinct functions in regulating collagen behaviors. The different ways in which matrix PGs interact with collagen have important implications for understanding the role of the ECM in diseases such as fibrosis and cancer, and suggest that matrix PGs are potential therapeutic targets.

**Highlights:**

- Small leucine rich proteoglycans (SLRPs) and large chondroitin sulfate (CS) proteoglycans (PGs) have distinct effects on collagen fibrous network behavior.
- Unlike other matrix proteoglycans, versican promotes collagen fibrillogenesis in an *in vitro* spectrophotometric (turbidity) assay.
- The versican core protein has a larger impact on collagen behavior in a fibrillogenesis assay than its glycosaminoglycan chains do.
- Versican increases the diameter of collagen fibers and the porosity of collagen fibrous networks, unlike aggrecan and SLRPs.
- The addition of versican to collagen does not alter fibroblast contractility but leads to enhanced cell-mediated collagen reorganization and contraction.

## Introduction

The compositional and structural complexity of the extracellular matrix (ECM) is important for maintaining appropriate cell and tissue function [1]. The ECM consists of a 3D network of fibers, primarily type I and other fibrillar collagens, in the form of cross-linked fibrous networks. The structure and organization of these networks can be regulated by cell-generated forces [2] and by interactions with other ECM components including proteoglycans (PGs) and glycosaminoglycans (GAGs) [3][4]. PGs are highly negatively charged (especially the large PGs with multiple GAG side chains) and, in part through their interactions with collagen and water, contribute to tissue mechanics by swelling and stiffening tissues, which enables them to resist compression and to retain water [5]. They are also important regulators of ECM related diseases including inflammation, fibrosis and cancer [6][7].

There are two groups of matrix (interstitial) PGs. The first is the family of small leucine-rich PGs (SLRPs), which have core proteins of about 50-60 kDa attached to 1-4 GAG chains; this group includes decorin, biglycan, fibromodulin, lumican and others. SLRPs have been well studied as collagen regulators [8][9], and the binding sites between collagen and some of the SLRPs have been identified through a combination of crystal structures and solid-phase binding data [10][11]. SLRPs are crucial for regulating collagen fiber formation and organization during development, especially in tissues such as cornea and tendon that require a highly-organized collagen network for their functions. Biglycan- and lumican-deficient mice show a disrupted lamellar structure in the cornea and impaired corneal transparency [12], and decorin-, fibromodulin- and lumican-deficient mice have tendons with irregular fiber morphology, abnormal fiber diameter distributions, and atypically non-uniform interfibrillar spaces [13][14].

The second family of matrix PGs is the hyalectan family of large chondroitin sulfate (CS) PGs, which includes versican (with a core protein of approximately 360 kDa and 12-15 CS chains) and aggrecan (with a core protein of approximately 250 kDa and around 100 GAG chains, including both CS and keratan sulfate (KS)) [15][16]. Compared with SLRPs, large PGs have a significantly larger mass of negatively-charged GAG side chains (with total molecular weights of 1-2.5 MDa) and they can bind hyaluronic acid (HA) to form even larger space-occupying aggregates [17][18]. Versican is universally distributed throughout the human body and has roles in regulating tissue morphogenesis and homeostasis and in the matrix response to injury [19][20], while aggrecan is predominantly expressed in cartilage and blood vessels [21]. Versican has at least 5 different isoforms that are generated by alternative splicing, and that have different distributions, degrees of GAG modification, and potentially functions [22][23]. Aggrecan interacts with collagen through its KS binding domain, as shown by a solid-phase binding assay [24]. There is one report, also based on a solid-phase binding result, that versican binds to type I collagen [25], but the physical nature of the interaction between versican and collagen has not been well defined and the effects of large PGs on fibrillogenesis are overall not well understood. There is a particular need to clarify the role of versican in regulating collagen behavior given its widespread distribution and regulated expression.

We report here an investigation into the effects of matrix PGs, particularly versican, on collagen fibrous network behavior. We report that different matrix PGs (even within a particular family) have distinct roles in the regulation of collagen behavior, suggesting that the relative expression of individual matrix PGs may be an important regulator of tissue function and cell behavior in disease.

## Results

### Matrix proteoglycans have different effects on collagen fibrillogenesis *in vitro*

Proteoglycans and their GAG side chains have been well studied as collagen regulators [26] through the use of *in vitro* spectrophotometric (fibrillogenesis) assays whereby the turbidity of a collagen solution is measured as gelation proceeds [27], generating a sigmoidal curve with a lag phase followed by a growth phase and then, after complete gelation, a plateau. During the lateral growth of collagen fibrils, the formation of large aggregates contributes to increases in turbidity due to the increased molecular weight of the aggregates and alterations in particle scattering, reflected in absorbance at 400 nm [28]. While this assay does not directly measure fibrillogenesis, increments in turbidity reflect collagen fibril/fiber formation and changes in collagen organization; the assay has been widely used to identify factors that impact fibrillogenesis [28][29]. We tested both rat tail telocollagen and bovine atelocollagen in the *in vitro* collagen fibrillogenesis assay and studied the effect of versican in both cases (Fig. 1A). We found that gelation time for bovine atelocollagen was longer than for rat tail telocollagen. This is expected because it has been reported that telopeptides can function as docking sites, guiding collagen monomer alignment and lateral growth, and thus the diffusion time for collagen monomer addition to telocollagen would likely be lower than for atelocollagen [30]. Addition of versican isolated from bovine liver, which consisted primarily of the large, GAG-modified V0 and V1 isoforms, accelerated fibrillogenesis and increased the height of the plateau for both forms of collagen. Because of the rapidity of telocollagen gelation and our desire to study modulators of the process, we used atelocollagen for our remaining experiments, reasoning that it would better enable us to evaluate differences in collagen behavior with different matrix PGs added.

**Fig. 1.**
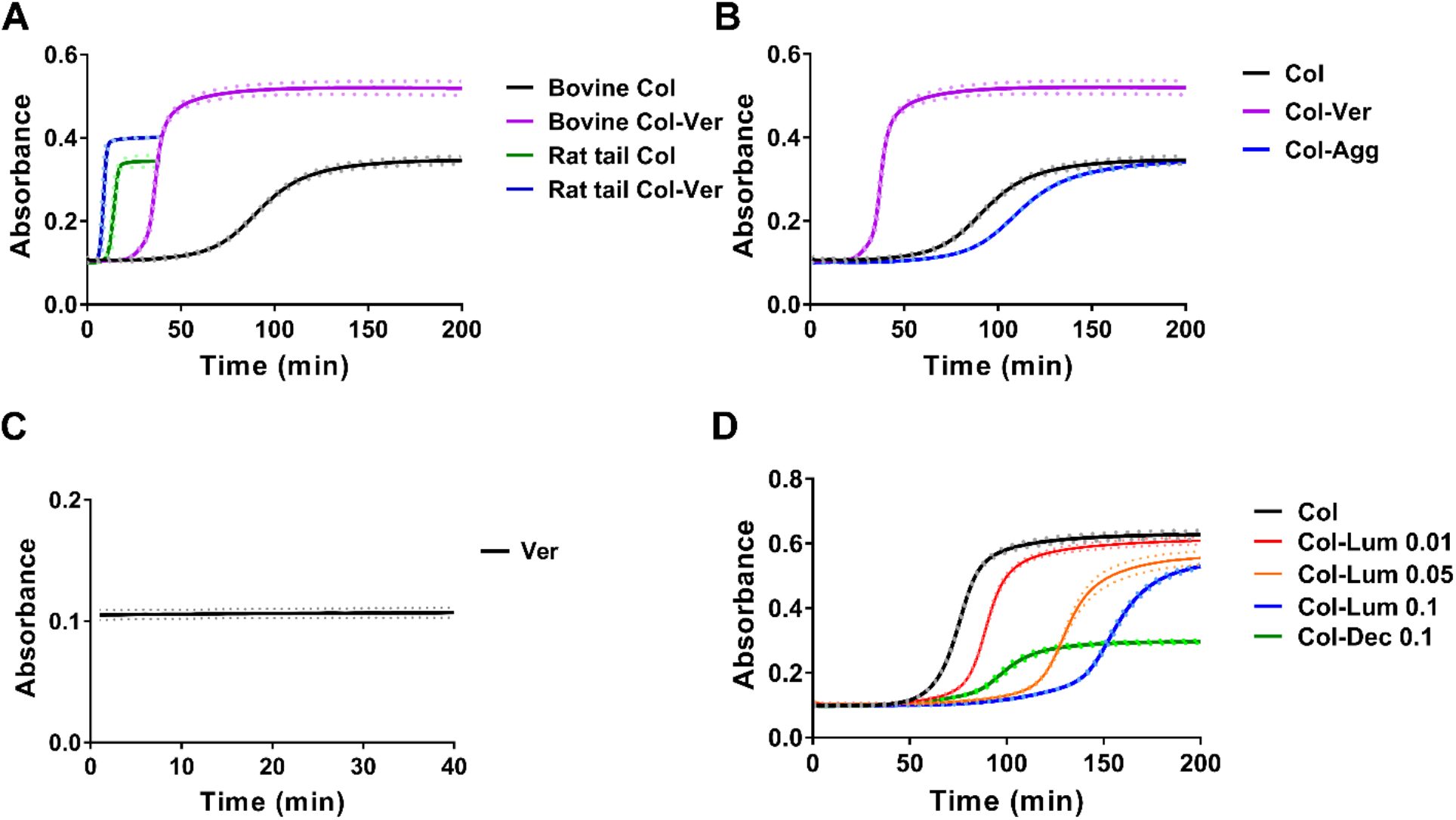
Different matrix proteoglycans have distinct effects on collagen fibrillogenesis in an *in vitro* turbidity assay. (A) Versican (Ver; 0.1mg/ml) was added to rat tail telocollagen (Col; 1.5 mg/ml) and bovine atelocollagen (1.5 mg/ml). (B) Versican (Ver; purple curve) or aggrecan (Agg; blue curve), both at 0.1 mg/ml, were added to atelocollagen (Col; 1.5mg/ml, black curve). Versican accelerated gelation dramatically while aggrecan slightly right-shifted the turbidity curve. (C) Versican alone (0.1 mg/ml) failed to gel and showed no change in turbidity over time under the assay conditions. (D) The SLRPs lumican (Lum; 0.01, 0.05 and 0.1 mg/ml) and decorin (Dec, 0.1 mg/ml) were added to atelocollagen (Col; 1.5 mg/ml). Decorin had a larger impact on decreasing fibrillogenesis than lumican. For all turbidity assays under all testing conditions, the pH and gelation temperature were the same. For all panels except C, three independent experiments were carried out for each condition, each with three technical replicates. Because there can be day-to-day differences in the absolute absorbance values for the assay, a representative figure from one experiment with mean curves is shown for each condition; however, all assays in a panel were carried out in parallel, and relative values among the different conditions were consistent in each individual experiment. The dotted lines represent Standard Deviation (SD). C was performed once with three technical replicates; the dotted lines represent SD.

To test the impact of versican versus aggrecan on collagen fibrillogenesis, we combined either of the two large PGs with atelocollagen before initiating the gelation assay. When versican was added to collagen, increases in turbidity of the mixture were more rapid and the plateau was higher than for collagen alone (Fig. 1B, 2C, purple and black curves). Carrying out the assay under identical conditions with versican alone showed no significant change in turbidity (Fig. 1C), suggesting that the change in the collagen curve with the addition of versican was due to interactions between versican and collagen. Surprisingly, the addition of the structurally-related large PG aggrecan to collagen slowed fibrillogenesis without changing the plateau (Fig. 1B, blue curve). The addition of CS (a GAG that modifies both versican and aggrecan) in unbound form (in a distribution of sizes ranging from 10-150 nm, as shown in Supplementary Fig. 1) left-shifted the turbidity curve in a dose-dependent manner, although less markedly than for intact versican even at similar concentrations (Fig. 2A). We then tested whether the versican protein core alone could alter collagen fibrillogenesis. We added the recombinant versican V3 isoform, which only contains the G1 and G3 domains, to telo- and atelocollagen and found it had the same effects on accelerating fibrillogenesis and increasing the plateau (Fig. 2B). We also removed the CS side chains by digesting with chondroitinase ABC (ChABC) (followed by dialysis against diH2O to remove the small, digested chains) and we observed an impact on the rate and plateau of the turbidity curves that was only slightly less than seen with the intact protein (Fig. 2C). In tests with enzyme-treated material, we confirmed that the heat-inactivated ChABC had minimal effect on this assay (Fig. 2C, blue curves). Thus, the versican core protein, with at best a minor contribution from its GAG side chains, modulates collagen gelation.

**Fig. 2.**
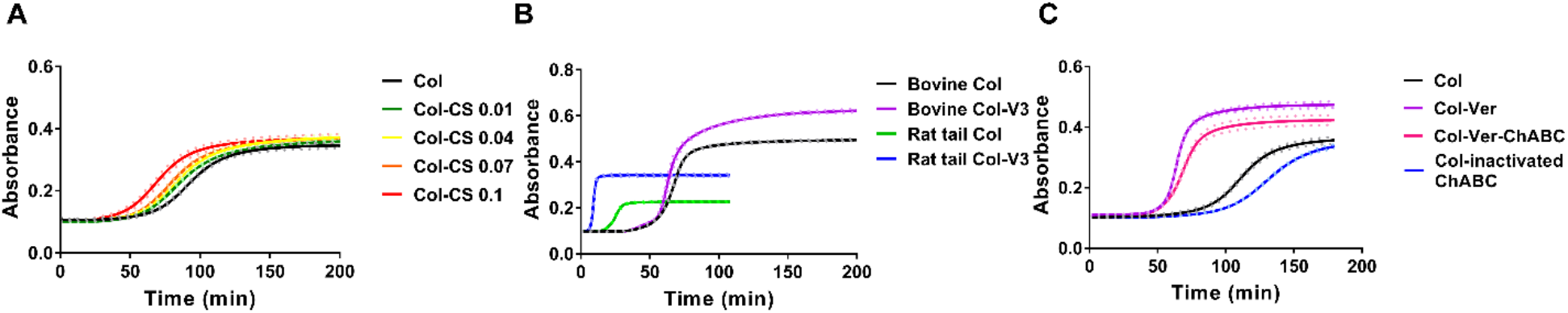
Versican core protein, with a minor contribution from the CS side chains, regulates collagen gelation. (A) Chondroitin sulfate (CS; 0.01, 0.04, 0.07 and 0.1 mg/ml; green, yellow, orange and red curves) was added to collagen (Col; 1.5 mg/ml; black curve). (B) Recombinant V3 isoform (V3, 0.1 mg/ml) was added to rat rail telocollagen (1.5 mg/ml) and bovine atelocollagen (1.5 mg/ml). (C) After digestion of the versican CS with ChABC, the remaining versican core protein was added at 0.1 mg/ml (pink curve) to atelocollagen (1.5 mg/ml; red curve) and caused a similar although slightly blunted right shift to the curves. Heat-inactivated ChABC had minimal effect on collagen gelation (blue curve). Three independent experiments were carried out for each condition, each with three technical replicates. Because there can be day-to-day differences in the absolute absorbance values for the assay, a representative figure from one experiment with mean curves is shown for each condition; however, all assays in a panel were carried out in parallel, and relative values among the different conditions were consistent in each individual experiment. The dotted lines represent SD.

The versican preparation we used was contaminated with a small amount of decorin (Supplementary Fig. 2), so we then tested whether the SLRPs had different effects on collagen fibrillogenesis in this assay and whether decorin could account for the effects noted using our versican preparation. We observed that the addition of both lumican (recombinant core protein with no GAGs) and decorin (full structure with GAGs, extracted from bovine articular cartilage) decreased the rate of collagen fibrillogenesis; decorin had particularly marked effects on both the rate and plateau and lumican decreased fibrillogenesis rate in a dose-dependent manner (Fig. 1C). The presence of decorin in the non-recombinant versican preparation is unlikely to account for the effects observed in the gelation assay (Fig. 1A, B) given that decorin alone had opposite effects (Fig. 1D) and that the recombinant form of versican (Fig. 2B) had similar effects as the form we isolated. Thus, we conclude from Figures 1 and 2 that matrix proteoglycans have different effects on collagen fibrillogenesis, regardless of their sizes and GAG modifications.

### Matrix proteoglycans have distinct effects on collagen fibrous networks

Scanning electron microscopy (SEM) of collagen matrices gelled plus or minus PGs was used to further analyze the impact of PGs on the collagen network. We tested both telo- and atelo-collagen (Supplementary Fig. 3). Telocollagen was used in this assay because the gelation of atelocollagen took up to 2-3 times as long (see Fig. 1A), raising concerns that dehydration might occur during atelocollagen solution gel formation. The addition of versican to collagen resulted in significantly enlarged fiber diameter and decreased porosity of the network as compared to collagen alone (Fig. 3A, B). The addition of aggrecan did not alter the diameter of collagen fibers or network porosity (Fig. 3C). The addition of decorin, but not lumican, decreased the diameter of collagen fibers slightly (Fig. 3D, F). Importantly, gelled samples were dehydrated as part of the preparation for SEM, causing the network to lose its native hydrated structure and the volume occupied by PGs due to negative charges to significantly decrease. The relative thickness of the fibers and porosity of the networks, however, are likely to persist.

**Fig. 3.**
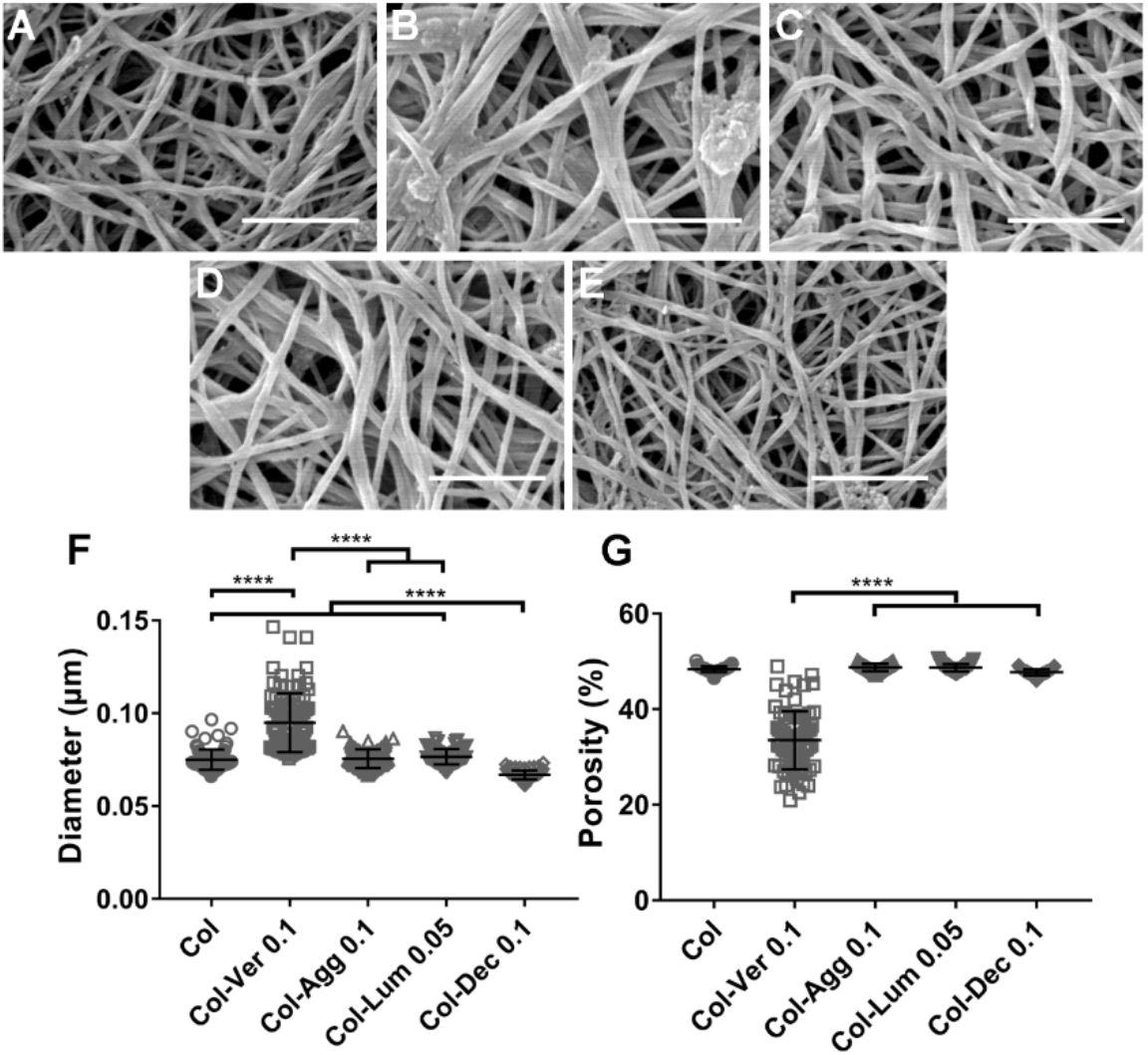
Matrix PGs have different effects on the structure of collagen networks. (A-E) Representative SEM images of telocollagen matrices with different PGs added. (A) Telocollagen (1.5 mg/ml) alone; (B-E) Telocollagen (Col; 1.5 mg/ml) with 0.1 mg/ml versican (Ver) (B), 0.1 mg/ml aggrecan (Agg) (C), 0.05 mg/ml lumican (Lum) (D) and 0.1 mg/ml decorin (Dec) (E). (F, G) Quantification of fiber diameter and porosity using DiameterJ. Three independent experiments were carried out and one gel was generated for each condition in each experiment. 5 SEM images were taken for each gel at random locations. When analyzed using FibrilTool, 5 sections were cropped from each SEM image and a measurement was taken on each cropped figure. Each data point represents a single measurement. Scale bar = 1 μm. Data represent mean ± SD. ****P<0.0001.

### Matrix proteoglycans regulate cell-mediated collagen compaction and alignment differently

The impact of PGs on cell-mediated collagen reorganization was studied using an *in vitro* model mimicking collagen organization and long-range force transmission at the tissue level [2]. In this assay, pairs of contractile cell spheroids (of either NIH 3T3 fibroblasts or primary liver portal fibroblasts) were placed atop collagen gel mixtures, and cell contractility-mediated collagen alignment and compaction were visualized using collagen second harmonic generation imaging (SHG) [31]. We mixed versican, aggrecan, decorin or lumican with collagen and allowed full gelation to occur, then placed fibroblast spheroids on the gels and imaged the collagen fibers after 24 hours of potential cell-mediated reorganization (Fig. 4A-C, blue). In this assay, increased SHG signal (blue) reflects increased local concentration and alignment of collagen. As was also shown in the *in vitro* turbidity assay, versican and aggrecan had distinct effects. The addition of versican, but not aggrecan, significantly increased cell-mediated collagen compaction (Fig. 4A-C, D), although there were no differences seen between any of the conditions in collagen organization in regions of the gels distant from cells. Interestingly, cell-mediated compaction of collagen in the collagen-versican mixture was sensitive to pH at pH values ranging from 7.20 to 7.40 (Fig. 4G). Cell-mediated compaction in the pure collagen plug, however, was not sensitive to pH in this range (Fig. 4F). There was no significant difference in anisotropy between any of the conditions, indicating that fibers in all conditions were equally parallel in the aligned area (Fig. 4E).

**Fig. 4.**
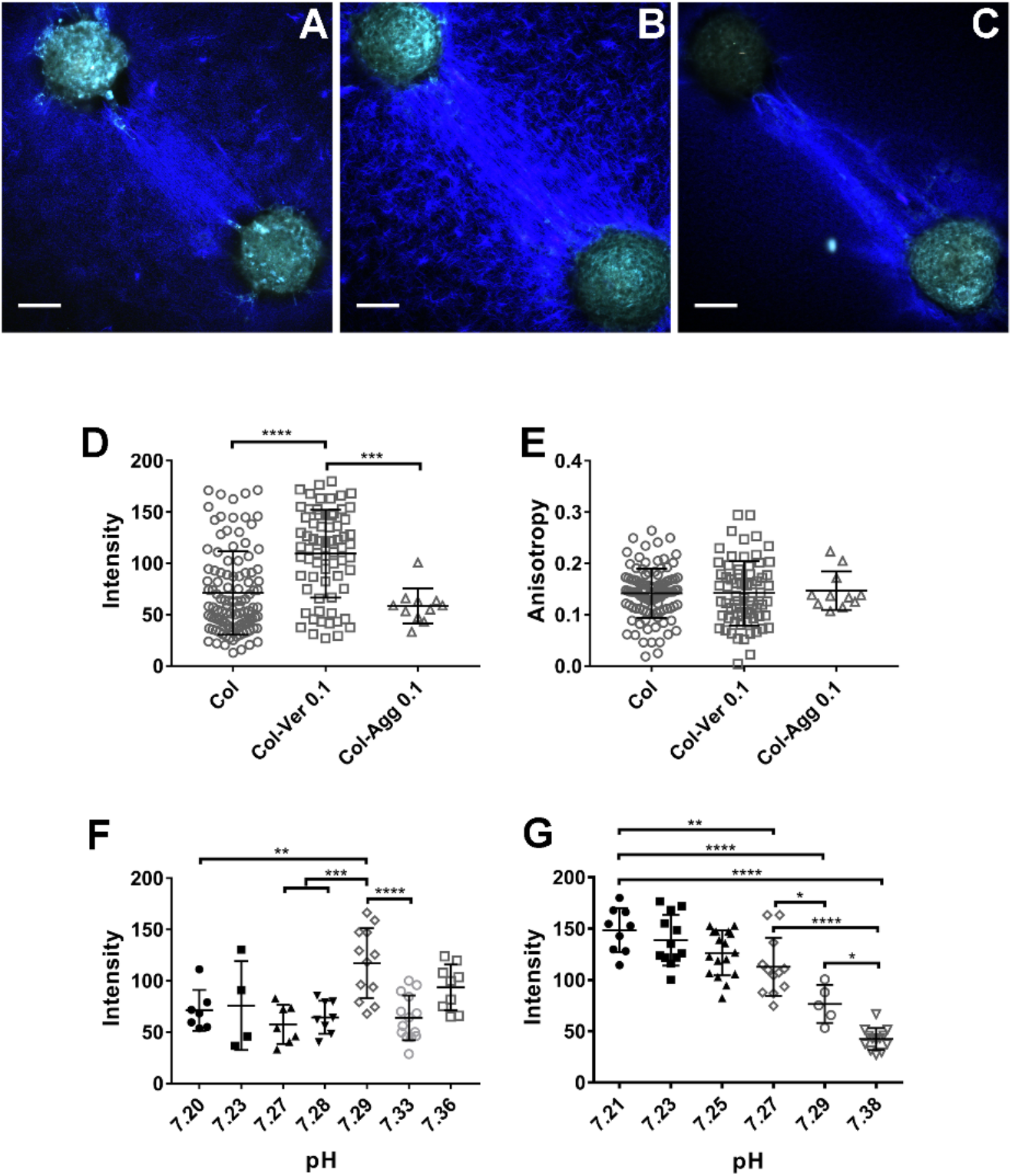
Large CS proteoglycans have differential effects on cell-mediated collagen reorganization. (A-C) Representative SHG images of aligned collagen fibers between pairs of NIH 3T3 spheroids. Blue represents the SHG signal from collagen; green is cell autofluorescence. (A) collagen (Col; 1.5 mg/ml) alone, (B) collagen-versican (Ver; 0.1 mg/ml) and (C) collagen-aggrecan (Agg; 0.1 mg/ml) plugs. (D-E) Intensity and anisotropy in the aligned collagen area for A-C. (F,G) Collagen compaction in pure collagen plugs (F) was not pH sensitive, but the impact of versican on collagen compaction was highly pH-dependent (G). Each data point in D-G represents collagen behavior between one pair of spheroids. At least 3 independent experiments were carried out for each condition, with at least 3 pairs of plugs examined for each experiment. For the pH testing in F and G, 4-12 pairs of spheroids were analyzed for each pH. Spheroids were seeded approximately 500 μm apart. Scale bars = 100 μm. Data represent mean ± SD. *P<0.05, **P<0.01, ***P<0.001 and ****P<0.0001.

We used spheroids of portal fibroblasts to assess the impact of decorin and lumican. There was a significant decrease in collagen compaction with the addition of either SLRP (Fig. 5A-D, E). Interestingly, the inclusion of decorin decreased changes in anisotropy significantly, although anisotropy was similar under all other conditions (Fig. 5F). To rule out changes in cell contractility on different matrices as an explanation for the observed differences in collagen compaction, traction force microscopy was used to measure contractility directly (Supplementary Fig. 4). We found that the addition of PGs to collagen did not lead to altered cell contractility (although with the caveat that this was done in 2D), suggesting that PGs regulate collagen behaviors through either direct or indirect effects on the structure of the fibrous network.

**Fig. 5.**
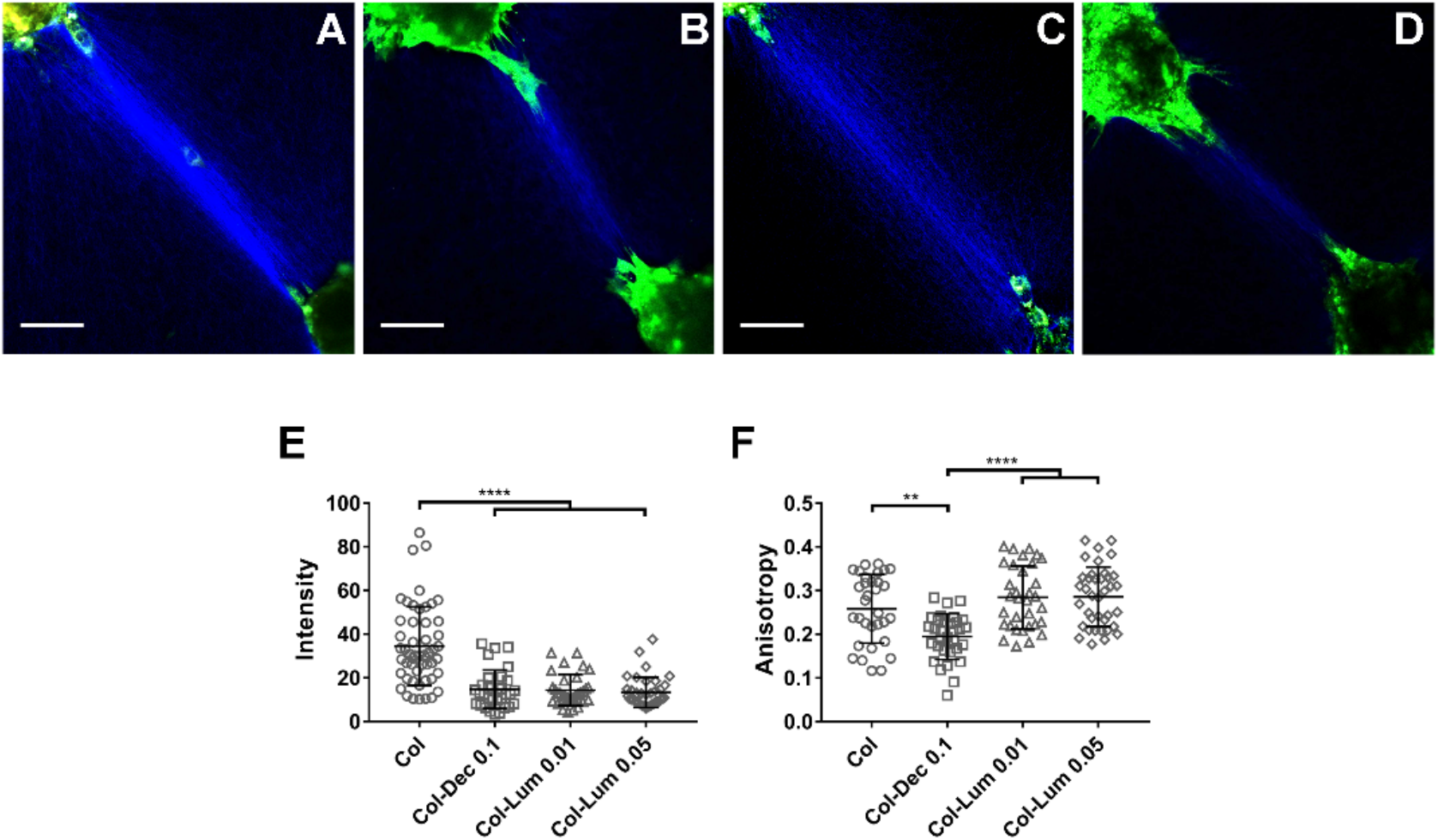
SLRPs regulate cell-mediated collagen reorganization differently. (A-D) Representative SHG images of collagen fibers between portal fibroblast spheroids on (A) collagen (1.5 mg/ml) alone, (B) collagen-decorin (Dec; 0.1 mg/ml) and (C, D) collagen-lumican (Lum, 0.01 or 0.05 mg/ml) plugs. (E, F) Quantification of cell-mediated collagen alignment with the addition of decorin and lumican, from A-D. Each data point in E and F represents collagen behavior between one pair of spheroids. At least 3 independent experiments were carried out for each condition, with at least 3 pairs of plugs in each experiment. Spheroids are seeded approximately 500 μm apart. Scale bar = 100 μm. Data represent mean ± SD. **P<0.01, ***P<0.001 and ****P<0.0001.

### Matrix proteoglycans have different roles in altering engineered microtissue contraction

We then used engineered microtissues (μTUGs) to determine whether the presence of PGs altered cell-mediated compaction of collagen matrices in 3D. Microtissues were generated by gelling collagen/cell mixtures in PDMS microwells with pairs of cantilevers; microtissue contractility resulted in displacement of the cantilevers (Fig. 6A, B). Representative light microscopic images of microtissues (pure collagen with NIH 3T3 cells, Fig. 6A, B) showed the displacement of the cantilevers in the presence and absence of microtissues. SHG imaging showed that the collagen fibers in engineered microtissues are well organized and aligned (Fig. 6C). Analysis of a large number of microtissues with and without PGs showed that the addition of versican significantly increased microtissue contraction while addition of aggrecan did not. For the SLRPs, decorin (0.1 mg/ml) and lumican (0.05 mg/ml) resulted in decreased contraction compared to collagen alone, while the addition of lumican at a lower concentration (0.01 mg/ml) had no effect. The addition of matrix PGs had no effect on fibroblast contractility tested by 2D traction force microscopy or on fibroblast proliferation culturing 24 h in contractile collagen gels (Supplementary Figs. 4, 5). Thus, we have shown that matrix PGs function as collagen regulators, with different effects on cell-mediated microtissue contraction.

**Fig. 6.**
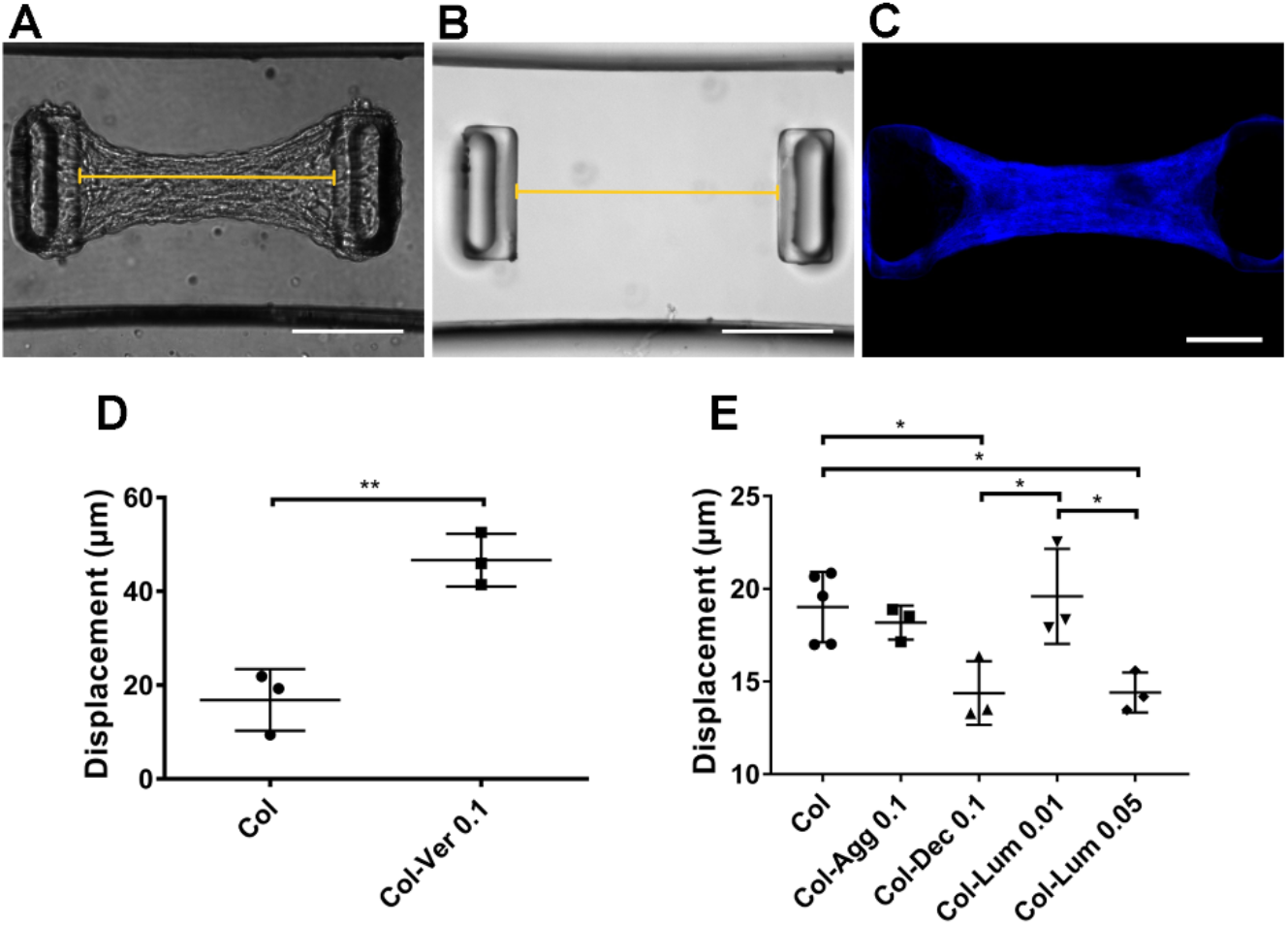
Matrix PGs have different effects on the contraction of engineered collagen microtissues. (A, B) Representative light microscopy images of PDMS cantilever displacement in μTUGs. (C) SHG imaging of μTUGs made using collagen and NIH 3T3 fibroblasts. (D) Quantification of increased displacement observed with inclusion of 0.1 mg/ml versican (Ver) in 1.5 mg/ml collagen (Col) microtissue. (E) Quantification of the displacement observed in collagen microtissues with or without aggrecan (Agg; 0.1 mg/ml), decorin (Dec; 0.1 mg/ml), or lumican (Lum; 0.01 mg/ml or 0.05 mg/ml). N>30 microtissues per each platform, three independent experiments (platforms) per condition. Points represent mean per platform. Scale bar = 200 μm. Data represent mean ± SE. *P<0.05 and **P<0.01.

## Discussion

Matrix PGs are important regulators of collagen fibrillogenesis and cell-mediated reorganization both *in vitro* and *in vivo*. We report here that different PGs, regardless of their structural similarity, have distinct effects on collagen behaviors. Versican, a widely-distributed hyalectan PG, has particularly notable behaviors compared to other PGs. It accelerates collagen gelation and upregulates cell-mediated collagen compaction and contraction, while aggrecan, another large hyalectan PG, slows gelation and has no effect on cell-mediated collagen reorganization. SLRPs, which also belong to the group of matrix PGs, similarly have opposing effects on collagen behaviors when compared to versican.

Previous *in vitro* spectrophotometric (fibrillogenesis) assays of collagen gelation have shown that decorin, lumican and biglycan (both the full protein and the core protein were tested) slow down fibrillogenesis, with a lower fibril formation plateau compared to collagen alone, and that their GAG side chains play a significant role in these effects [32][33][34]. Similarly, the use of atomic force microscopy to scan a mica disc coated with collagen-PG mixtures showed that adding the recombinant core proteins of decorin and lumican to collagen resulted in larger interfibrillar spaces and decreased fibril diameters [35]. We made similar observations using the *in vitro* spectrophotometric assay, although the differences in the kinetic curves for collagen with the addition of decorin are more pronounced (much flatter, with a lower plateau) than for those with the addition of lumican. One potential explanation for the difference between decorin and lumican is that decorin has 12 leucine rich repeats [36] and lumican has 11 [37], resulting in different geometries of their otherwise similar horseshoe-shaped, leucine-rich protein domains. These domains, which bind collagen at the C- and N-terminal domains [38], may have different effects on collagen spacing during fibrillogenesis. Decorin, for example, interacts with charged residues in the d band of the collagen α1 chain via the charged residues on its inner surface [10], and may do so differently than lumican. Another possible explanation relates to the source of SLRPs we used. The decorin used was a native PG isolated from bovine cartilage, and has one CS or DS side chain close to its N-terminus, while the lumican we used was recombinant, without the native GAG chains (*in vivo*, 4 KS chains on the leucine-rich domain) [39][40]. The negatively-charged side chain in decorin might cause physical repulsion that inhibits fibril lateral growth and thus, the different types, numbers and locations of side chains on SLRPs may cause distinct effects on the interactions between collagen and SLRPs which could affect fibrillogenesis.

In contrast to SLRPs, large matrix PGs have not been well studied as modulators of collagen fibrillogenesis. This is particularly true for versican, which, unlike the more widely-studied aggrecan, is distributed throughout the body. Published work has focused primarily on studying the functional role of versican on cell and tissue phenotype in development and disease [41][42][43]. Interestingly, versican and aggrecan, in spite of significant structural similarities, have distinct effects in multiple assays, as reported here. We observed that versican accelerates collagen gelation while aggrecan had a modest negative effect. Our in vitro findings on aggrecan are consistent with recently reported in vivo data suggesting that the loss of aggrecan led to enhanced surface fibrillation in cartilage [44]. A typical aggrecan chain has approximately 100 GAG chains; physical repulsion caused by negatively-charged GAGs bound to collagen potentially limits the interactions between collagen fibrils, leading to slowed fibrillogenesis. Versican, in contrast to aggrecan, has only about 10-15 GAG chains, although they are all CS (which is longer and contains more negatively charges than KS) rather than a mixture of CS and KS. Our data suggest that the core protein of versican rather than its GAG chains is the main determinant of its effects on collagen. The versican core protein may bind collagen more tightly than the aggrecan core protein, or the GAG chains (potentially the KS chains found on aggrecan but not versican) may play a role in aggrecan interactions with collagen. CS has a complicated and controversial role in regulating fibrillogenesis. While some published work [45][46] shows that CS chains increase the rate of fibrillogenesis, other work suggests the opposite [47]. Although our findings suggest that CS slightly accelerates collagen gelation, we also find that the concentration of the CS chains has an impact.

SEM imaging of collagen matrices provided detailed visualized and quantitative data on the impact of matrix PGs on the structure of the collagen network, but in a dehydrated state. It has been reported that the presence of versican, aggrecan or the SLRPs (mainly decorin) decreases collagen fibril width [26]. Our data show that versican and aggrecan, which are both large CS PGs, have different effects on the structure of the collagen network. The results highlight the potentially unique role of versican on collagen fibrils and the network. Evidence addressing the regulation of SLRPs on the structure of the collagen network is contradictory. Raspanti et al. [48] showed that the presence of decorin in a 1:5 ratio with collagen promoted collagen fusion and increased collagen fibril diameter. Reese et al., however, found that the inclusion of decorin into collagen matrices at a ratio of 1:40 resulted in a denser network with thinner fibrils [34]. Our data show that addition of decorin at a ratio of 1:15 with collagen yields thinner fibrils in a looser network, suggesting a potential dose dependence that needs to be further investigated. For lumican, Rada et al. found that its addition into the collagen network *in vitro* resulted in thinner collagen fibrils, as visualized by transmission electron microscopy [33], while *in vivo* studies by Chakravarti et al. showed that the diameter of collagen in corneal stroma was increased in lumican-deficient mice [49]. Our data suggest that lumican has minimal effect on collagen fiber diameter, but there are no other SEM data published for comparison.

Contractile cells can generate force, which acts on ECM fibers and mediates ECM organization [2][50]; structural and mechanical stimuli from the ECM can also feedback on cells and impact cell behavior through conversion into biochemical signals. This reciprocal crosstalk between cells and the ECM is important in regulating cell function and tissue morphogenesis. The collagen plug assay enables the study of both cellular and matrix factors regulating cell-mediated long-range force transmission, which is one manifestation of this crosstalk. This assay is of particular interest because it may serve as a model of *in vivo* pathology such as the bridging fibrosis typical of advanced liver fibrosis [51]. SLRPs including decorin and lumican decrease collagen fiber compaction, while versican (but not aggrecan), increase collagen condensation. The finding (using traction force microscopy) that fibroblast contractility is not altered when PGs are added to collagen in 2D suggests that the behavior of the collagen network is altered by specific interactions between PGs and collagen fibers and not by differences in cell behaviors. The arch-like shape of decorin can occupy the space around collagen to limit parallel fibril assembly via binding with collagen α1 chain [34], which is consistent with our observation that the addition of decorin blunted the increase in anisotropy of collagen fibers in response to cell contractility. Decorin, as a structural spacer, would make it harder for contractile forces to stretch fibers closer in a linear fashion.

Higher level tissue contraction is also mediated by collagen organization. Engineered microtissue gauges represent a recently developed technique to investigate both cell contractility and ECM contraction in 3D. This technique has been used to study the organization of matrix proteins (including collagen and fibronectin) in response to applied force or cell contractility [52]. It has been shown previously that the presence of decorin in collagen gels or culture media inhibits collagen gel contraction, which is consistent with our results [53][54]. Another report suggested that the inclusion of lumican at very small amount (approximately 0.4 ng/ml) increased fibroblast-mediated collagen gel contraction; in contrast, we found that 10 ng/ml (consistent with the concentration in native tissues [55]) had no effect on contraction, while the inclusion of lumican at 50 ng/ml decreased microtissue contraction. Understanding the potentially dose-dependent effect of lumican on collagen behavior will require further investigation. Versican has been shown to upregulate fibroblast-mediated collagen gel contraction [56], which is the same behavior we find in our microtissues. However, aggrecan has no effect on microtissue contraction. As matrix PGs have no effects on cell contractility in 2D and cell proliferation after 24h culturing in contractile collagen gels (shown in Supplementary Figs. 3, 4), we conclude that the distinct roles of PGs in altering tissue contraction are due mainly to their different effects on the structure and organization of the collagen network.

In sum, we observe distinct effects of different matrix PGs, even from the same subfamily, on collagen fibrillogenesis and the organization of collagen fibrous networks. Interestingly, versican appears to enhance fibrillogenesis while aggrecan and the SLRPs have the opposite effect. This suggests that the precisely-controlled deposition and the relative amounts of different PGs expressed in normal and disease states, including during development and in fibrosis and cancer, may have an important impact on collagen organization and on cell-ECM cross talk. Specific matrix PGs may be potential therapeutic targets; by controlling and altering their expression, it might be possible to control collagen behavior, including cell-generated collagen fiber reorganization, tissue contraction, and long-range cell-cell communication.

## Acknowledgements

This work was supported by the UPenn National Science Foundation MRSEC (DMR-1720530), by the Center for Engineering MechanoBiology (CEMB), a National Science Foundation Science and Technology Center (under grant agreement CMMI:15-48571), and by grant R01 EB017753 (to RGW). We are grateful to Gordon Ruthel and the Penn Vet imaging Core for extensive assistance with SHG imaging; Yuri Veklich and the University of Pennsylvania Cell and Developmental Biology Microscopy Core for help with SEM; and the Image Analysis and Gene Expression Quantification Core of the National Institute of Diabetes, Digestive and Kidney Diseases Center for Molecular Studies in Digestive and Liver Diseases (NIH P30 DK050306). The versican antibody 12C5 developed by R.A. Asher was obtained from the Developmental Studies Hybridoma Bank, created by the NICHD of the NIH and maintained at The University of Iowa, Department of Biology, Iowa City, IA 52242.

## Experimental procedures

### Reagents, antibodies and cells

Bovine type I atelocollagen (lacking N- and C-terminal telopeptide regions) was from Advanced Biomatrix (San Diego, CA, USA) and rat tail type I telocollagen (with intact telopeptide regions) was from Corning (Corning, NY, USA). Versican was isolated from bovine liver as described below. Aggrecan was isolated from bovine cartilage [57]. Decorin from bovine articular cartilage was from Sigma (St. Louis, MO, USA) and human recombinant lumican protein (lacking GAG chains) was from R&D Systems (Minneapolis, MN, USA). CS sodium salt isolated from bovine cartilage and ChABC from *Proteus vulgaris* were from Sigma. Sylgard 184 PDMS and its curing agent were from Dow Corning (Midland, MI, USA). Trichloro silane, isopropanol, pluronic F127 and Medium 199 were purchased from Sigma; sodium bicarbonate from Corning; and CellPURE™ HEPES from Fisher Scientific (Hampton, NH, USA). Protease complete tablets were from Roche (Roche, Basel, Switzerland). 40% acrylamide and 2% bisacrylamide stock solutions were purchased from Bio-Rad (Bio-Rad Laboratories, Hercules, CA, USA). Tetramethylethylene diamine (TEMED), ammonium persulfate (APS) and a solution of 0.2 μm fluorescent beads was from Fisher Scientific. Coverslip activation reagents were aminopropyltrimethoxysilane (Sigma) and glutaraldehyde (Sigma). PAA gel surface activation reagents were ethyl(dimethylaminopropyl) carbodiimide (EDC) and N-hydroxysuccinimide (NHS) solution (Fisher Scientific). Collagenase from Clostridium histolyticum was purchased from Sigma.

Anti-versican antibody 12C5 was from DSHB (Developmental Studies Hybridoma Bank, Iowa city, IA, USA), anti-aggrecan antibody BC-3 was from Thermo Scientific and anti-decorin antibody ab175404 was from Sigma.

NIH 3T3 fibroblasts (CRL-1658™) were obtained from the ATCC^®^ (Manassas, VA, USA) and portal fibroblasts were isolated from rat liver as described [58]. Both types of fibroblasts were cultured in DMEM (Dulbecco’s Modification of Eagle’s Medium with 4.5 g/L glucose and L-glutamine without sodium pyruvate (Corning)) with 10% fetal bovine serum (Gemini Bio-Products, West Sacramento, CA, USA) supplemented with 1% penicillin/streptomycin (Corning) and 0.5% fungizone (Life Technologies, Carlsbad, CA, USA) at 37°C in a humidified atmosphere with 5% CO_2_/balance air.

### Dynamic light scattering

A dynamic light scattering nano-sizer (Zetasizer, Malvern, Westborough, MA, USA) was used to quantify the size of the CS sodium salt from Sigma, which was isolated from bovine cartilage. CS was diluted to 0.01 mg/ml with PBS and 0.5 ml CS solution was loaded into a glass cuvette. The cuvette was inserted into the instrument and the number of measurements was set to 3.

### Versican isolation

Versican was isolated from bovine liver by a modification of a published protocol [59]. Briefly, bovine liver was mechanically disrupted and treated with extraction buffer containing 4 M guanidine hydrochloride, 100 mM sodium acetate and protease complete tablets (Roche, Basel, Switzerland) (pH=7.2) at 4°C for 72 h. Tissue residue was removed by centrifugation for 1 h at 16,000×g. Cesium chloride was added to the supernatant solution until the density reached 1.59 g/ml and then spun at 100,000×g for 24 h. 1 ml fractions were taken carefully from the top to the bottom and the density of each fraction was measured. Fractions above a density of 1.54 g/ml were dialyzed against 1 M sodium chloride for 24 h and against diH2O for 24 h. Samples were re-concentrated using a 100k centrifugal filter (Millipore Sigma, Burlington, MA, USA). The composition of isolated samples and the presence of versican were confirmed by dot blotting using anti-versican antibody 12C5, anti-aggrecan antibody BC-3 and anti-decorin antibody ab175404 (Supplementary Fig.2).

### In vitro spectrometric (turbidity) assay

Type I bovine atelocollagen was diluted to a final concentration of 1.5 mg/ml. All solutions were kept on ice before gelation. Briefly, 187.5 μL collagen solution (3.2 mg/ml) was gently mixed with 40 μL 10× PBS, 4 μL 1N NaOH, and 168.5 μL deionized water (diH2O). In some cases, type I rat tail telocollagen was used and prepared similarly. For some experiments, versican, aggrecan and decorin were added to the collagen solution to a final concentration of 0.1 mg/ml; lumican was added to 0.01 and 0.05 mg/ml. CS side chains were tested by adding CS at 0.01, 0.04, 0.07 and 0.1 mg/ml. The versican core protein was obtained by treating the intact protein with 250 mU chondroitinase ABC (ChABC) per mg substrate (in 50 mM sodium acetate, pH=8.0) at 37°C overnight, followed by dialysis against distilled water to remove the small CS chains and confirmation of the absence of CS chains using the Blyscan assay (Bicolor, UK). The pH of all collagen solutions was carefully adjusted to 7.4; solutions were incubated on ice for exactly 1 h before pipetting into 96-well plates. The absorbance of the solution was read at 400 nm by a plate reader (Infinite 200 Pro, Tecan Life Sciences) at 37°C until gelation was complete (when the absorbance curve reached its plateau) [26]. For all gelation assay experiments, all conditions compared in a given graph were tested at the same time.

### Scanning electron microscopy

Rat tail type I telocollagen was diluted to a final concentration of 1.5 mg/ml and supplemented with different PGs as descried above for the spectrometric (turbidity) assay. It was polymerized on 8 mm coverslips for 25 min. The bovine atelocollagen gel sample was made similarly. The collagen gels were fixed with 2.5% glutaraldehyde in cacodylate buffer overnight at 4°C. The samples were further processed by the Cell and Developmental Biology Microscopy Core (University of Pennsylvania, Philadelphia, PA, USA). Briefly, samples were dehydrated with a graded series of ethanol washes (50, 75, 90, 95, 100%) and incubated with 50% hexamethyldisilazane (HDMS) for 30 min. Samples were then incubated with 100% HDMS three times and air dried before mounting on stubs. Samples were imaged on a FEI Quanta 250 FEG scanning electron microscope (Thermo Scientific). Bovine atelocollagen was prepared and studied in the same manner. 5 SEM images were taken per each gel at 5 random locations with 10,000 × magnitude. 5 randomly-cropped figures (384×256 pixels) from each SEM image were analyzed using DiameterJ, an imageJ plugin, which was used to quantify fiber diameter and network porosity (porosity is the area of pores over the total area of the figure) [60].

### Collagen plug assay

Rat tail type I collagen was diluted to a final concentration of 1.5 mg/ml with 10× PBS and diH2O as for the gelation assay. Versican, aggrecan or decorin were added to the collagen solution to concentrations of 0.1 mg/ml; lumican was added to 0.01 and 0.05 mg/ml. The pH of the collagen solution was adjusted to 7.4 and incubated on ice for 1 h before pipetting into a microwell plate with a glass-bottomed cutout (14 mm Microwell, MatTek, Ashland, MA). The plate was sealed with parafilm and kept in an incubator (5% CO_2_/balance air) overnight at 37°C. Fibroblast spheroids were formed by the hanging droplet method [61]. Briefly, cells were trypsinized and suspended in DMEM at 25,000 cells/ml (for NIH 3T3 cells) and 200,000 cells/ml (for portal fibroblasts). 20 μL droplets were placed on the underside of a petri dish lid. To avoid drying, 10 ml DMEM was added to the dish. After inversion of the lid, the cell droplets were cultured for 5 days (for NIH 3T3) or 3 days (for portal fibroblasts). At the time of seeding, 1 ml media was added on top of each collagen gel. Spheroids were captured by a 20 μL pipette and carefully placed on the gel in pairs approximately 500 μm apart. After spheroids were cultured on gels for 24 h, gels were fixed with 10% formalin for 10 min. SHG imaging using a Leica SP5 spectral imaging confocal/dual-photon microscope was used to image collagen fiber reorganization between each pair of spheroids [62]. Aligned collagen fibers were analyzed using Image J. The pixel intensity of each aligned region was quantified and the FibrilTool plug-in [63] was used to define the anisotropy of alignment in selected bridging areas.

### Engineered microtissue assay

Micro-tissue gauges (μTUGs) were fabricated as per a published protocol. Briefly, the mold was rinsed with isopropanol, plasma coated and salinized in a vacuum chamber overnight. PDMS was mixed with its curing agent (10:1) for 5 min and degassed. PDMS was pipetted on top of the stamps and again degassed. PDMS was also placed on 35 mm petri dishes to cover the bottom and incubated at 65°C for 30 min. After degassing, the stamps were inverted and placed in the center of dishes. The dishes were then filled with PDMS and incubated at 65°C overnight. After incubation, stamps were removed and the μTUG platforms were rinsed with ethanol and isopropanol. 1.5 mg/ml collagen solution was prepared as in Table 1. Versican, aggrecan and decorin were added to reach a final concentration of 0.1 mg/ml while lumican was used at 0.01 and 0.05 mg/ml. The pH was adjusted to 7.4 and the collagen solution was incubated on ice. The platforms were sterilized with UV light for 15 min and rinsed with 70% ethanol, then rinsed with 0.2% pluronic F127 and centrifuged at 500×g until there were no bubbles in the wells. After rinsing the platforms twice with PBS, 1 ml collagen-PG solution was added to each and degassed for 3 min. The platforms were then centrifuged at 700×g for 2 min and stored at 4°C for avoiding gelation. NIH 3T3 fibroblasts were harvested from culture plates and 150,000 cells were mixed with 0.5 ml collagen solution before gently pipetting to mix, then added to each platform. The platforms were spun at 206×g for 2 min, then turned 90 degrees and spun for another 2 min. Extra solution was carefully aspirated and the platforms were placed inverted in a centrifuge and spun at 37×g for 15-20 s. 1 ml PBS was added to the lid and the platforms were incubated at 37°C for 20 min until gelation. 1.5 ml culture media was added to each platform and the platforms were incubated at 37°C for 24 h, until microtissues had formed. Images were taken using a light microscope (Leica DM IRM) before and after removal of microtissues by pipetting and rinsing with PBS. Cantilever displacements were measured by ImageJ and used to determine microtissue contraction.

**Table 1.** Components of collagen gels in engineered microtissue assay

|  |  |
| --- | --- |
| diH<sub>2</sub>O | 1067 μL |
| M199 (10×) | 200 μL |
| HEPES (250mM) | 80 μL |
| NaHCO<sub>3</sub> (5% w/v) | 14 μL |
| NaOH (1M) | 24 μL |
| Collagen (4.88 mg/ml) | 615 μL |

### Traction force microscopy

Traction force microscopy was used to study the effect of matrix PGs on cell contractility. The protocol was modified from previous publications [64][65]. Briefly, 7.9 kPa polyacrylamide gels (Table 2) were made by mixing 40% acrylamide and 2% bisacrylamide stock solutions with tetramethylethylene diamine (TEMED) and 1% ammonium persulfate (APS). This gel solution was mixed with 0.2 μm fluorescent bead solution (diluted at 1:1000) and then covered with a 25 mm glass coverslip pre-activated with 0.5% aminopropyltrimethoxysilane and 0.5% glutaraldehyde. After polymerization for 30 min, the gel surface was activated with ethyl(dimethylaminopropyl) carbodiimide (EDC) and N-hydroxysuccinimide (NHS) solution (17.5mg/ml NHS and 10 mg/ml EDC in milliQ water) and coated with collagen mixed with different matrix PGs and either cellular or plasma fibronectin. 3T3 Fibroblasts were seeded at 20,000 cells per gel and incubated overnight. Live cell imaging was applied using EVOS AUTO2 (Thermo Invitrogen) and single cell images were taken before and after removing cells with 10% sodium dodecyl sulfate. The average traction force was calculated by measuring the displacement of fluorescent beads (ImageJ plugin available at https://sites.google.com/site/qingzongtseng/tfm) [66].

**Table 2.** The protocol for making 7.9 kPa polyacrylamide gel

| Stiffness (Pa) | 40% acrylamide (μL) | 2% bisacrylamide (μL) | 10×PBS (μL) | diH<sub>2</sub>O (μL) | TEMED (μL) | 1% APS (μL) | Total volume (μL) |
| --- | --- | --- | --- | --- | --- | --- | --- |
| 7900 | 187.5 | 35 | 100 | 576.5 | 1 | 100 | 1000 |

### Fibroblast proliferation in contractile collagen gels

The same numbers of 3T3 fibroblasts used in the μTUG assays were cultured in contractile collagen gels for 24h and cell proliferation was measured. Briefly, after preparing collagen solutions with different PGs, NIH 3T3 fibroblasts were mixed with gel solution and incubated at 37°C for 20 min. Then, gels were carefully detached from each well and cultured for 24 h. The contractile collagen gels were digested with 10 mg/ml collagenase for 15 min; the cell numbers were counted and compared among each condition.

### Statistical analysis

All results were analyzed by GraphPad Prism 7 (San Diego, CA, USA) using unpaired t test or one-way ANOVA. P values were determined by Tukey’s multiple comparison test, in which *P<0.05 was considered to be statistically significant.

## Supplementary figures

**Supplementary Fig. 1.**
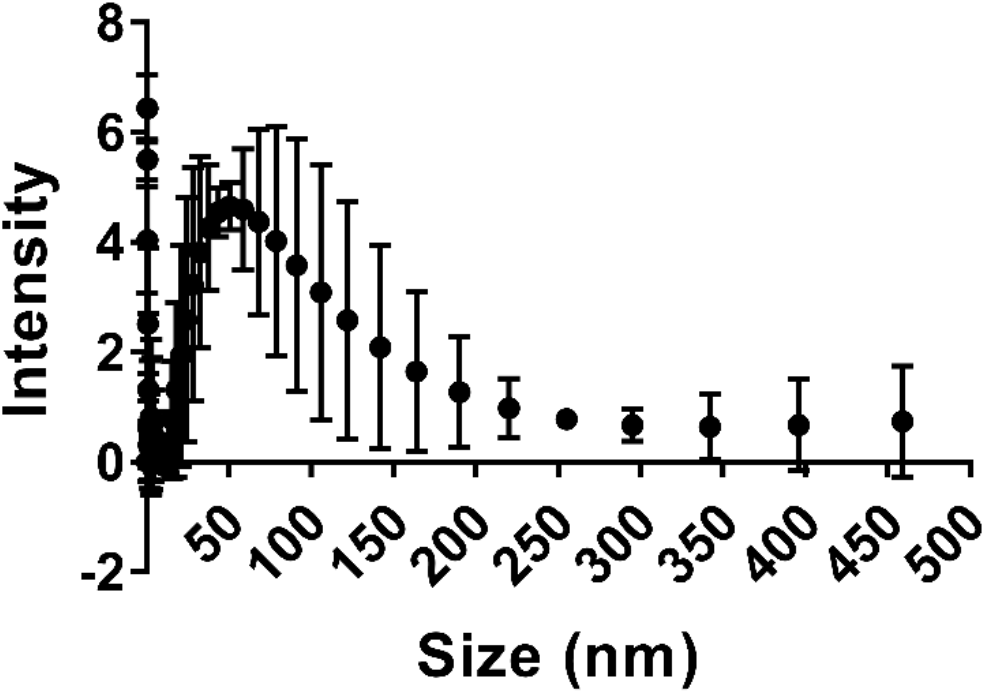
The size distribution of chondroitin sulfate tested by dynamic light scattering. CS sodium salt isolated from bovine cartilage (Sigma) has a distribution of sizes ranging from 10-150 nm.

**Supplementary Fig. 2.**
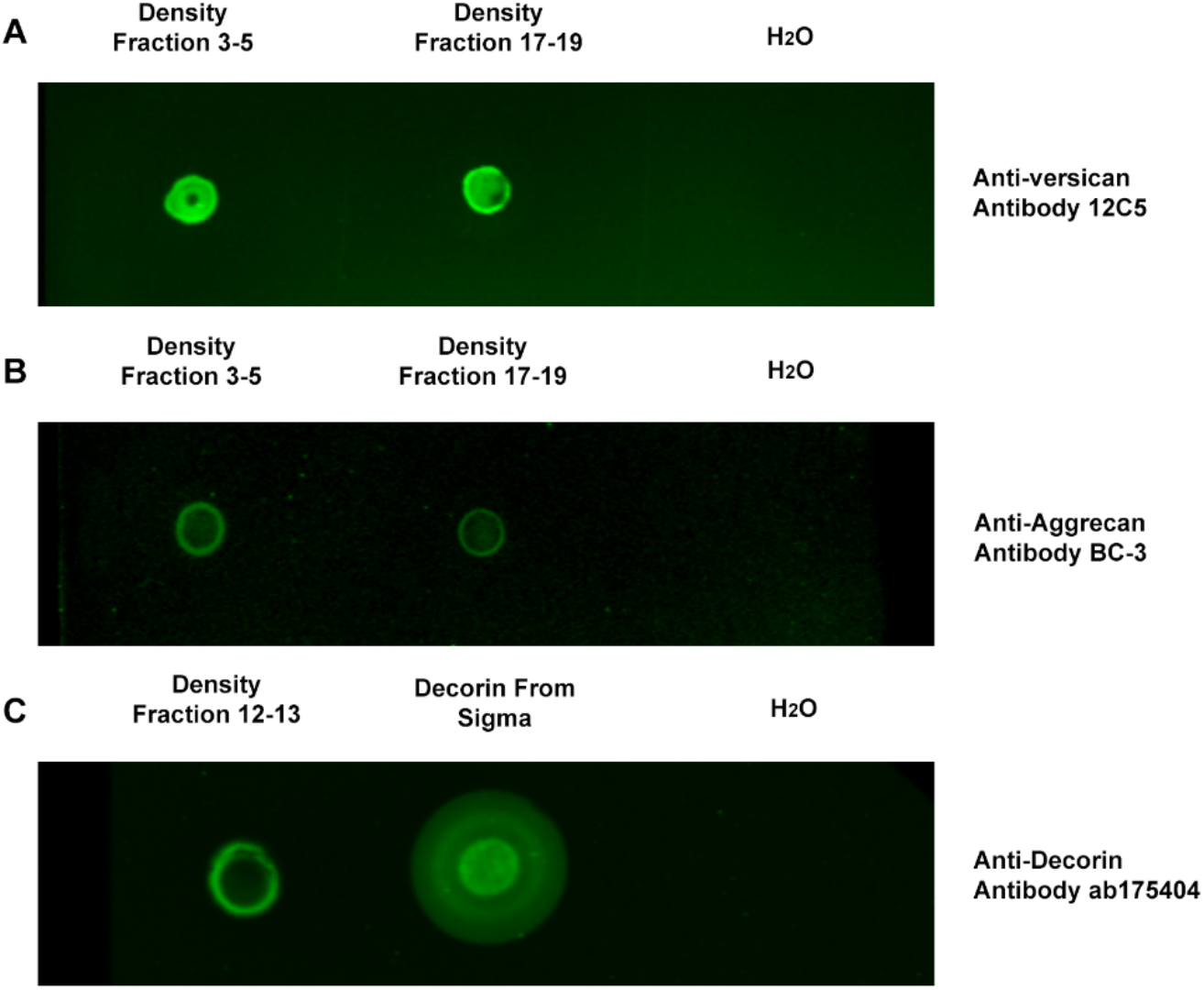
Purity of versican preparation. (A) Dot blotting column fractions and staining with versican antibody 12C5 confirmed the presence of versican. (B, C) Dot blots also demonstrated minor contamination of the versican sample with aggrecan (B) and decorin (approximately 0.37 mg/ml decorin in 4.68 mg/ml extracted sample) (C).

**Supplementary Fig. 3.**
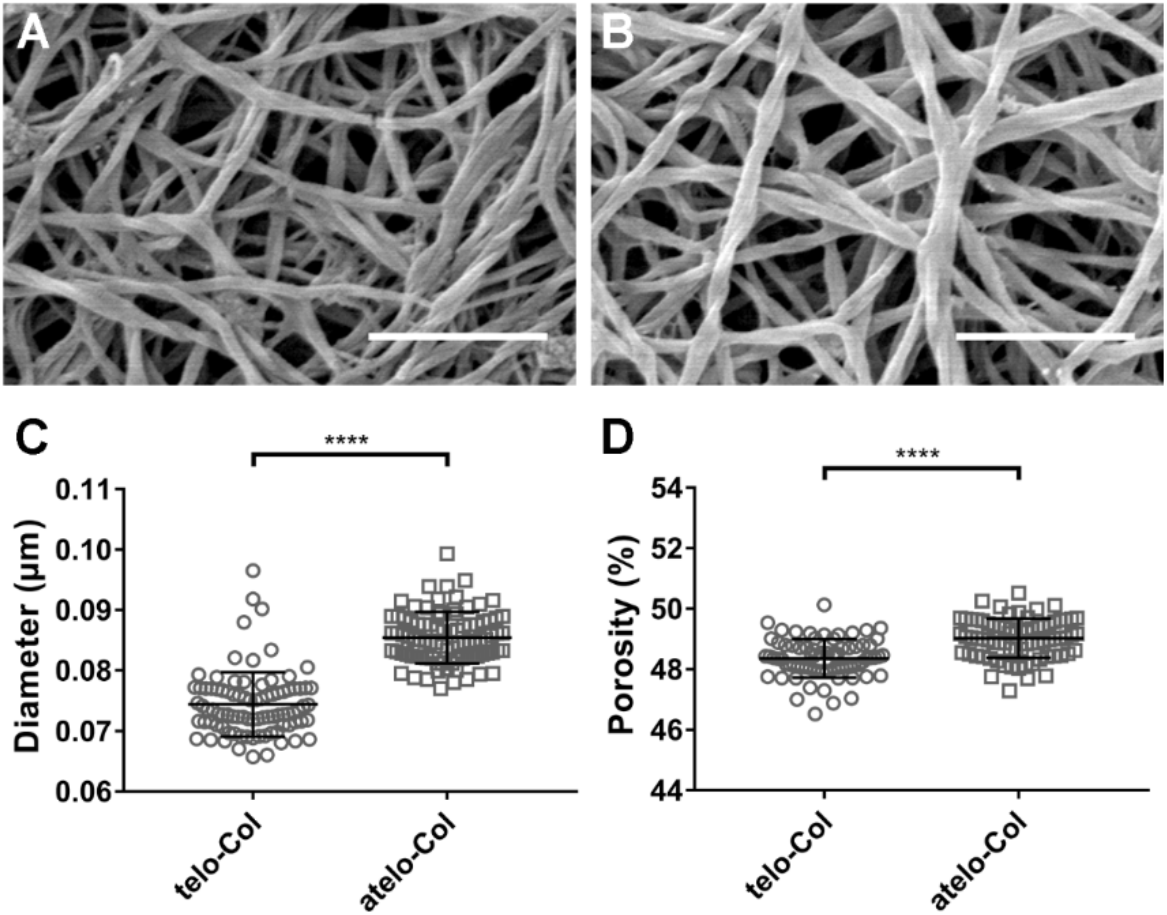
The structure of the telo- and atelo-collagen networks with SEM. Atelocollagen lacks the telo-peptide regions which are the most common sites of covalent crosslinks, and therefore not surprisingly formed a looser network. It also shows thicker fibers compared to telo-collagen. (A, B) SEM imaging of collagen matrices: (A) 1.5 mg/ml telocollagen; (B) 1.5 mg/ml atelocollagen. (C, D) Quantification of fiber diameter and porosity analyzed using DiameterJ. The data show thicker fibers and a looser network for atelocollagen. Three independent experiments were carried out for each condition. 5 images were taken for each gel and measurements taken on 5 random locations for each image. Scale bar = 1 μm. Data represent mean ± SD. ****P<0.0001.

**Supplementary Fig. 4.**
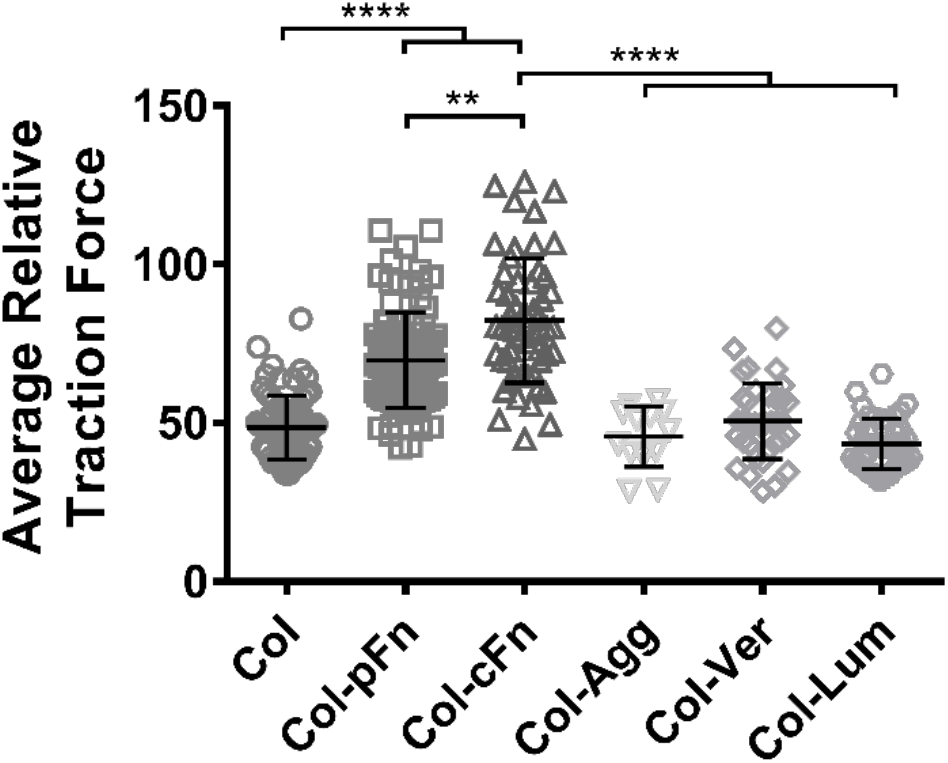
Traction force microscopy of NIH 3T3 cells on matrices of various composition. 7.9 kPa polyacrylamide gels were coated with 0.1 mg/ml collagen (Col) mixed with plasma fibronectin (pFn), cellular fibronectin (cFn), aggrecan (Agg), versican (Ver) and lumican (Lum) at 0.1mg/ml. The inclusion of PGs did not alter cellular contractility. In contrast, there was a significant increase with both variants of fibronectin, which are included for comparison. Three independent experiments were carried out for each condition. Data represent mean ± SD. **P<0.01 and ****P<0.0001.

**Supplementary Fig. 5.**
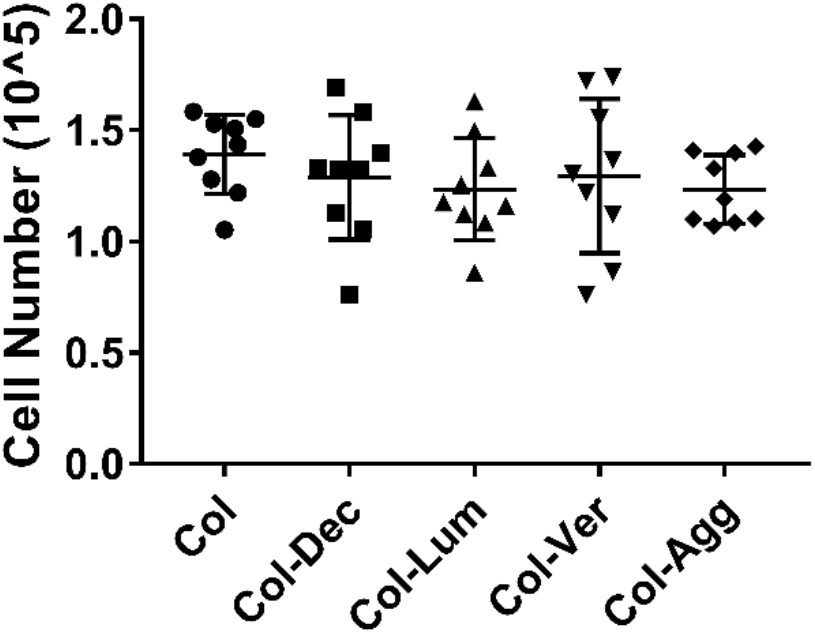
3T3 fibroblasts were cultured in contractile collagen gels with the same proteoglycan manipulations used in the μTUG assay (lumican was tested at 0.05 mg/ml); see Fig. 6. After culture for 24 h, collagen gels were digested with collagenase and the total cell number was counted. We confirmed that the matrix PGs we studied had no influence on proliferation of fibroblasts cultured in collagen gels when compared with collagen alone over 24 h. Three independent experiments were carried out for each condition.

